# Lifecycle dominates the volatilome character of the dimorphic fungus *Coccidioides* spp

**DOI:** 10.1101/2021.01.15.426916

**Authors:** Emily A. Higgins Keppler, Heather L. Mead, Bridget M. Barker, Heather D. Bean

## Abstract

Valley fever (coccidioidomycosis) is an endemic fungal pneumonia of the North and South American deserts. The causative agents of Valley fever are the dimorphic fungi *Coccidioides immitis* and *C. posadasii*, which grow as mycelia in the environment and spherules within the lungs of vulnerable hosts. The current diagnostics for Valley fever are severely lacking due to poor sensitivity and invasiveness, contributing to a 23-day median time-to-diagnosis, and therefore new diagnostic tools are needed. We are working toward the development of a breath-based diagnostic for coccidioidomycosis, and in this initial study we characterized the volatile metabolomes (or volatilomes) of *in vitro* cultures of *Coccidioides*. Using solid-phase microextraction and comprehensive two-dimensional gas chromatography coupled to time-of-flight mass spectrometry (GC×GC–TOFMS), we characterized the VOCs produced by six strains of each species during mycelial or spherule growth. We detected a total of 353 VOCs that were at least two-fold more abundant in a *Coccidioides* culture versus medium controls and found the volatile metabolome of *Coccidioides* is more dependent on growth phase (spherule versus mycelia) than on the species. The volatile profiles of *C. immitis* and *C. posadasii* have strong similarities, indicating that a single suite of Valley fever breath biomarkers can be developed to detect both species.

**IMPORTANCE:** Coccidioidomycosis, or Valley fever, causes up to 30% of community-acquired pneumonias in endemic and highly populated areas of the US desert southwest. The infection is difficult to diagnose by standard serological and histopathological methods, which delays an appropriate treatment. Therefore, we are working toward the development of breath-based diagnostics for Valley fever. In this study, we characterized the volatile metabolomes of six strains each of *Coccidioides immitis* and *C. posadasii*, the dimorphic fungal species that cause Valley fever. By analyzing the volatilomes during the two modes of growth for the fungus—mycelia and spherules—we observed that the lifecycle plays a significant role in the volatiles produced by *Coccidioides*. In contrast, we observed no significant differences in the *C. immitis* versus *C. posadasii* volatilomes. These data suggest that lifecycle, rather than species, should guide the selection of putative biomarkers for a Valley fever breath test.

## INTRODUCTION

Coccidioidomycosis, or Valley fever, is a disease that is endemic to the deserts of the western United States, Mexico, and Central and South America and is responsible for an estimated 350,000 new infections per year (1). In endemic and highly populated areas of the US, e.g., Phoenix and Tucson, Arizona and the San Joaquin Valley in California, up to 30% of community-acquired pneumonias may be caused by Valley fever, and with growing populations in these regions, Valley fever cases in the US are expected to climb (2). The causative agent of Valley fever is the opportunistic pathogen *Coccidioides*, a genus of dimorphic fungus comprised of two species, *Coccidioides immitis* and *C. posadasii*. In the environment, *Coccidioides* spp. grow in the soil during rainy periods as mycelia, which become dormant and form arthroconidia during dry periods. As the soil is disturbed, the arthroconidia can become airborne, and when inhaled, the fungus can initiate a parasitic lifecycle as spherules within the lungs of a susceptible host, causing pneumonia. The median duration of human Valley fever pneumonia is 120 days, costing $93,734 per person in direct and indirect costs (2, 3). Approximately three quarters of the diagnosed cases of Valley fever require antifungal therapy (4), and 1% of those infected will experience disseminated disease, with a third of those being fatal (4).

A definitive diagnosis for Valley fever requires a positive fungal culture or histological examination of tissue or bodily fluid. However, due to the invasiveness of obtaining suitable lung specimens for culture or histology, serological testing for antibodies against *Coccidioides* is the most commonly performed diagnostic test. Multiple serological tests are available for IgM, IgG, or complement-fixing antibodies, as well as an enzyme-linked immunoassay that uses proprietary coccidioidal antigens. However, these tests lack sensitivity for coccidioidomycosis (5, 6). Due to the difficulty in diagnosing Valley fever, it is often mistaken for bacterial pneumonia and inappropriately treated with antibiotics (7–10). The lack of a suitable diagnostic strongly contributes to an unacceptably long 23-day median time-to-diagnosis (2).

Breath-based diagnostics are a promising novel approach for diagnosing respiratory infections. Numerous *in vitro* studies have demonstrated that bacterial and fungal respiratory pathogens have unique volatile metabolome (or *volatilome*) signatures that can be used to differentiate and identify them to the genus, species, and strain level (11). Thus far, relatively few studies have specifically focused on identifying breath biomarkers for fungal lung diseases. *In vitro* analyses of the volatiles produced by *Aspergillus* spp. have shown that fungi produce many compounds that are also frequently detected in bacterial cultures, e.g., (alcohols) ethanol, propanol, (ketones) acetone, 2-nonanone, 2-undecanone, 2,3-butanedione, (sulfides) dimethyl disulfide, dimethyl trisulfide, and (pyrazines) 2-methyl pyrazine, and 2,5-dimethyl pyrazine (12–16). However, *Aspergillus* also produces a wider variety of monoterpenes, sesquiterpenes, and terpenoids, which are less common in bacterial volatilomes (14, 17). With this knowledge, Koo et al. used a biologically guided approach to identify volatile biomarkers of pulmonary invasive aspergillosis (IA), focusing on terpenes and terpenoids produced *in vitro* as putative biomarkers for a breath test (17). They determined that the *A. fumigatus in vitro* volatilome contained a distinctive combination of monoterpenes and sesquiterpenes, and in breath the sesquiterpenes were the most specific for differentiating patients with IA versus other pulmonary infections (17). This example for *A. fumigatus* establishes a proof-of-concept that direct detection of fungal metabolites in breath can be used as a noninvasive, species-specific approach to identify the underlying microbial cause of pneumonia.

We are taking an *in vitro*-guided approach toward developing a Valley fever breath test by first building a catalog of volatile compounds produced by *C. immitis* and *C. posadasii*. In this study, we have cultured six strains of each species and used solid-phase microextraction and comprehensive two-dimensional gas chromatography – time-of-flight mass spectrometry (GC×GC–TOFMS) to characterize the volatile organic compounds (VOCs) produced during mycelial or spherule growth. We expected to find significant differences in the VOCs between species and between lifecycles, but also posited that there would be a subset of VOCs that are common to *Coccidioides* spp. during spherule growth, which will serve as our candidate biomarkers for breath test development.

## RESULTS

### *Coccidioides* isolates in this study

The *Coccidioides* genus is widely distributed throughout North and South America. The genus is composed of two geographically (18) and genetically distinct species (18, 19), *C. posadasii* and *C. immitis*, which are further divided into five well-defined populations (20–22). *C. posadasii* has a more extensive biogeographic distribution and is divided into three populations, occurring in Texas/Venezuela, Mexico/South America, and Arizona. The species *C. immitis* is divided into two populations found in central and southern California, and recently Washington State, representing a newly discovered third distinct population (23, 24). Representative members of all populations cause disease in humans and other mammals (20–22). Due to the magnitude of observed genetic variation and geographic distribution between species, we hypothesized that metabolomic variation exists. To capture diversity across the volatilome between and within species, we included a total of twelve isolates from both species, representing four populations, all of which were isolated from humans (Table 1).

**Table 1.** *Coccidioides* spp. strains used in this study. All isolates are from human infections.

| Species | Strain | Population* | ATCC | NCBI** |
| --- | --- | --- | --- | --- |
| *C. posadasii* | Silveira | AZ | NR-48944 | ABAI000000000.2 |
|  | B3221 | AZ |  | NA |
|  | B3222 | AZ |  | NA |
|  | RMSCC2343 | TX/MEX/SA |  | SRR3468064 |
|  | RMSCC3506 | TX/MEX/SA |  | SRR3468053 |
|  | GT-166 | TX/MEX/SA |  | NA |
| *C. immitis* | RS | SDMX | NR-48942 | AAEC000000000.3 |
|  | RMSCC2395 | SDMX | NR-48938 | NA |
|  | RMSCC3505 | SDMX |  | NA |
|  | RMSCC2006 | SJV | NR-48934 | NA |
|  | RMSCC2009 | SJV |  | SRR3468015 |
|  | RMSCC2010 | SJV | NR-48935 | NA |

\*Populations are defined as *C. posadasii* collected from Arizona (AZ), *C. posadasii* collected from Texas/Mexico/South America (TX/MEX/SA), *C. immitis* collected from San Diego, California, and Mexico (SDMX), and *C. immitis* collected from San Joaquin Valley, California (SJV), as previously published (25). \*\*Published genotype data, if available (21).

### Characteristics of the *Coccidioides* volatilome by species and lifecycle

We cultured each *Coccidioides* isolate in biological triplicate using temperatures and oxygen concentrations that induce mycelial or spherule growth, but using the same culture medium, facilitating direct comparisons between the VOCs produced during the two lifecycles. Using solid-phase microextraction to adsorb VOCs from the headspace of fungal culture filtrates, and GC×GC–TOFMS to analyze the volatilomes, we detected a total of 353 chromatographic peaks that were at least two-fold more abundant in a *Coccidioides* culture (n = 72) versus the medium controls (n = 6) (Table S1). Of the 353 volatiles detected in *Coccidioides* cultures, 28 were assigned putative names based on mass spectral and chromatographic data, which included 13 compounds previously associated with human and environmental fungal pathogens (26) (Table 2). For the unnamed compounds, we assigned chemical classifications to 45 of them based on a combination of mass spectral and chromatographic characteristics (Table S1).

**Table 2.**
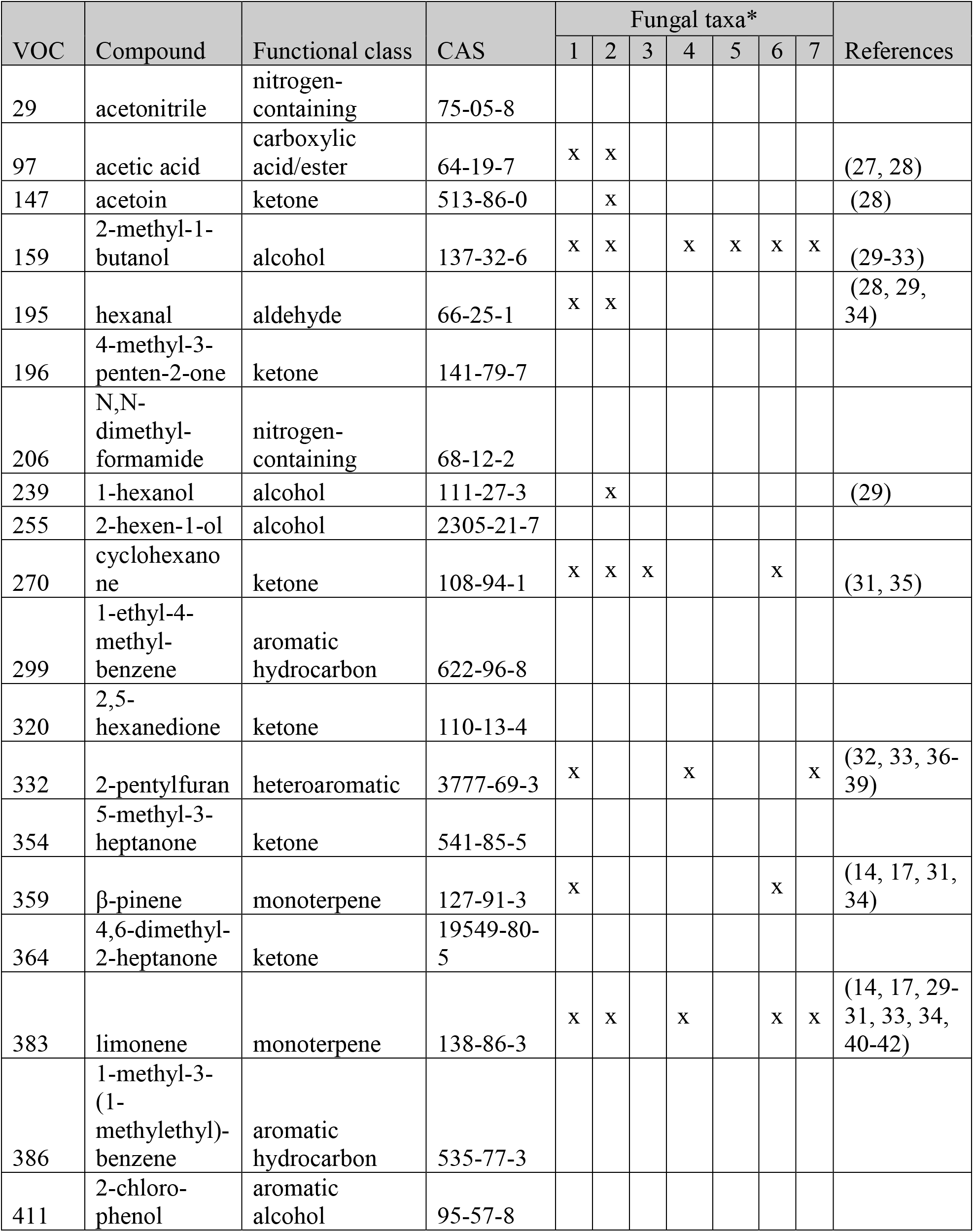
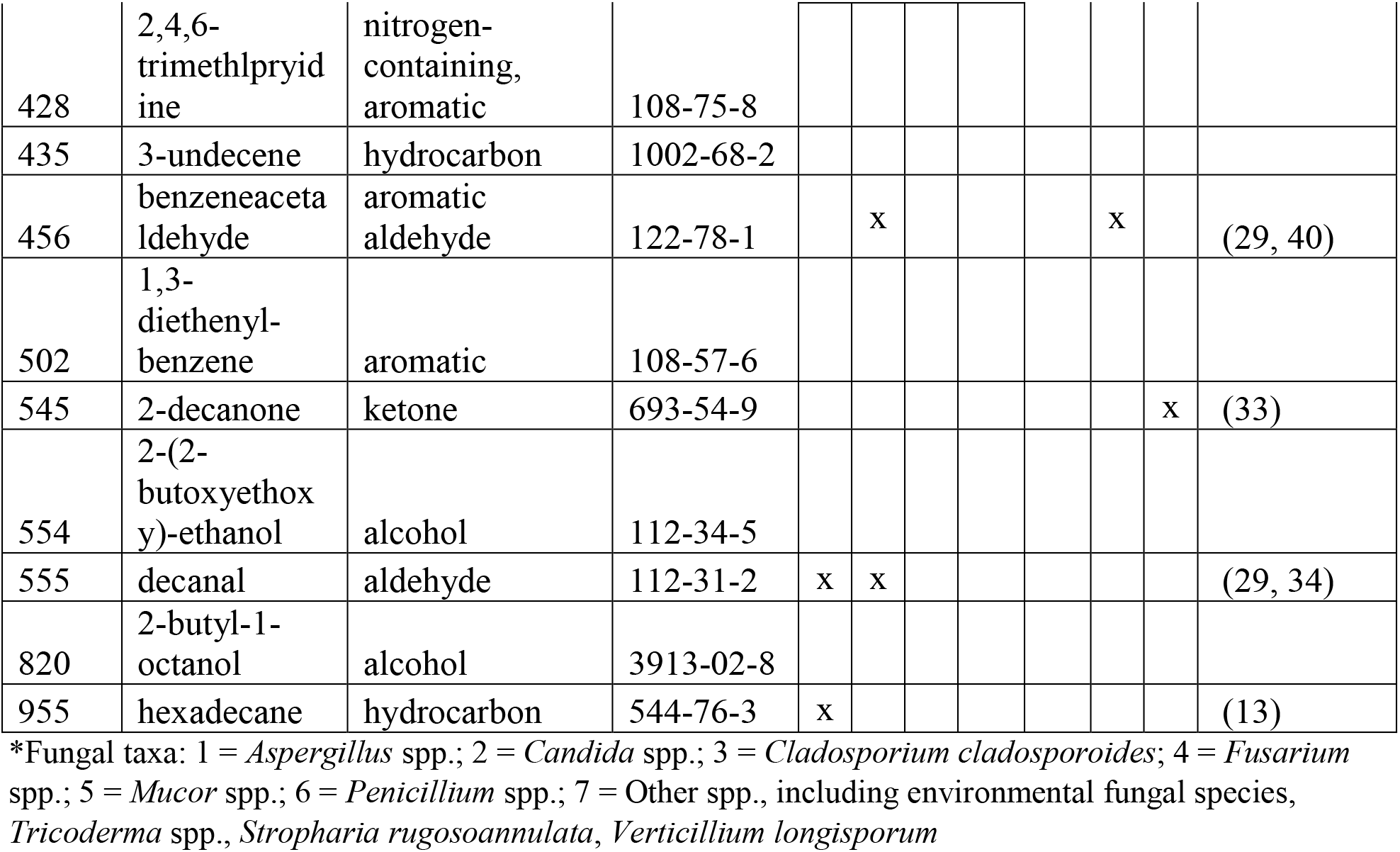
Named *Coccidioides* spp. volatiles and their previous reports in other fungal taxa.

The total number of volatiles produced by *C. posadasii* versus *C. immitis* were similar, at 291 versus 309, respectively (Figure 1; Table S2). Within each species, the two lifecycles are quite divergent in their volatilomes, sharing less than a third of the total volatilome (Figure 1 A, and B). However, within each lifecycle, the two species share many similarities, with more than half of the VOCs produced by both *C. posadasii* and *C. immitis* (Figure 1 C, and D). In aggregate across the genus, or subdivided by species, the spherule volatilome is larger than the mycelial volatilome, but this appears to be driven by considerable differences in volatilome sizes observed in a few isolates (*C. posadasii* B3221, GT-166; *C. immitis* RS; Table S2).

**Figure 1.**
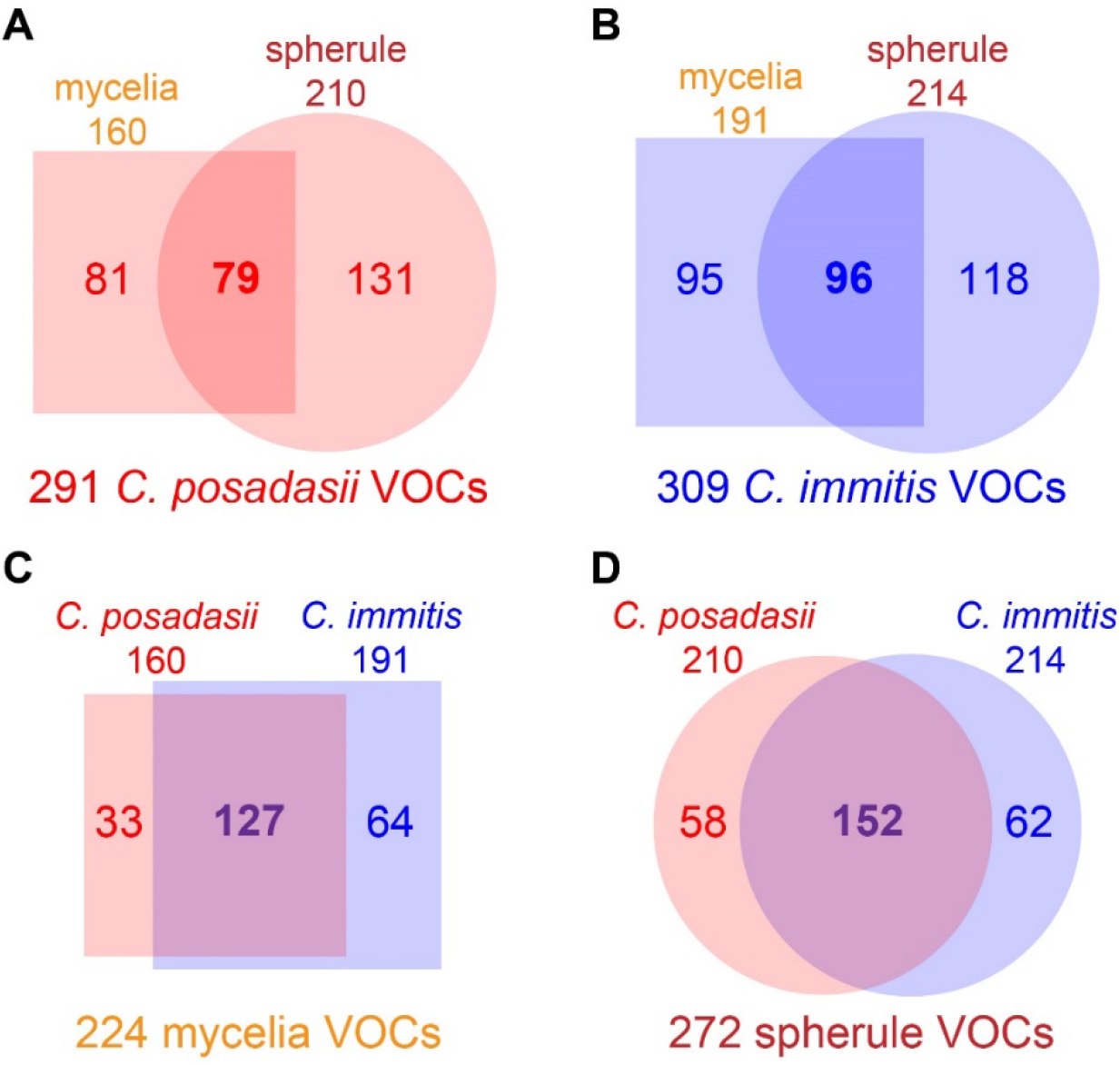
The numbers of analytes in at least two-fold greater abundance in a *Coccidioides* culture versus the media controls that are shared and unique among **A**) *C. posadasii* mycelia (square) vs. spherules (circle); **B**) *C. immitis* mycelia (square) vs. spherules (circle); **C**) mycelia of *C. posadasii* (red) vs. *C. immitis* (blue); and **D**) spherules of *C. posadasii* (red) vs. *C. immitis* (blue).

Analyzing the relative abundances of individual analytes, we observed some statistically-significant differences in VOCs produced during the two lifecycles, but not between the two species. We performed a Mann-Whitney *U*-test using Benjamini-Hochberg false discovery rate (FDR) correction comparing all spherule and mycelial VOCs, and identified 43 that were significantly different in abundance (p < 0.05) (Figure 2). Of the 43 volatiles, 32 of them were higher in abundance in spherule compared to 11 in mycelia, which mirrors the overall difference in the size of the volatilomes (Figure 1). Among the VOCs detected in higher abundance in one lifecycle versus another were 1-methyl-3-(1-methylethyl)-benzene and hexanal, which are associated with mycelia cultures, while 2-(2-butoxyethoxy)-ethanol, cyclohexanone, 2-hexen-1-ol, and 2,4,6-trimethlpryidine were associated with spherule cultures. There were no VOCs that were significantly different in relative abundance between *C. posadasii* and *C. immitis*.

**Figure 2.**
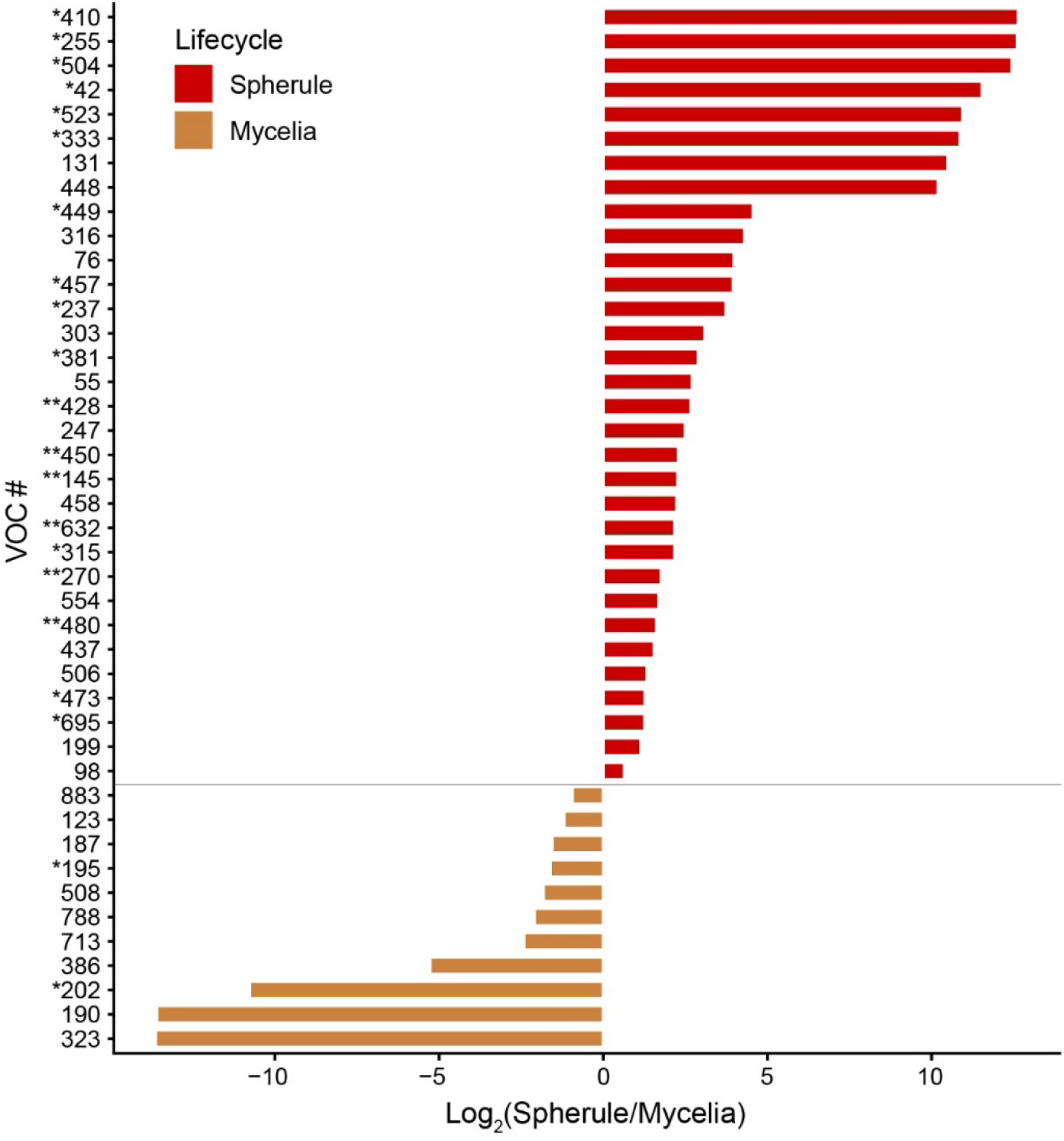
Forty-three volatiles significantly different in abundance between lifecycles (p < 0.05), expressed as log_2_ fold change in peak abundance. Maroon and tan represent spherule and mycelia peaks, respectively. Volatiles marked with an asterisks indicate greater statistically-significant differences (* p < 0.001, ** p < 0.0001). Compound IDs: 195 = hexanal; 255 = 2-hexen-1-ol; 270 = cyclohexanone; 386 = 1-methyl-3-(1-methylethyl)-benzene; 428 = 2,4,6-trimethlpryidine; 554 = 2-(2-butoxyethoxy)-ethanol.

Most (> 70%) of the spherule and mycelial VOCs detected in this study were produced by four or fewer *Coccidioides* strains in each lifecycle. Of the 272 spherule volatiles that were detected, only 35 (13%) were detected in at least two-thirds of all spherule cultures (Figure 3A), 12 of which were also significantly more abundant in spherule versus mycelia cultures. Nineteen of 224 mycelia volatiles (8%) were detected in at least two-thirds of all mycelia samples (Figure 3B), and only two were significantly different between lifecycles (VOC 199 and 428). However, both volatiles were produced in higher relative abundance in spherules. Only two VOCs, 39 and 748, were detected in two-thirds of both spherule and mycelia samples. Interestingly, the relative abundance of a VOC was usually consistent across all strains. For example, in every spherule culture in which limonene (VOC 383) was detected, it was at a relative concentration between two and tenfold greater than media (Figure 3A).

**Figure 3.**
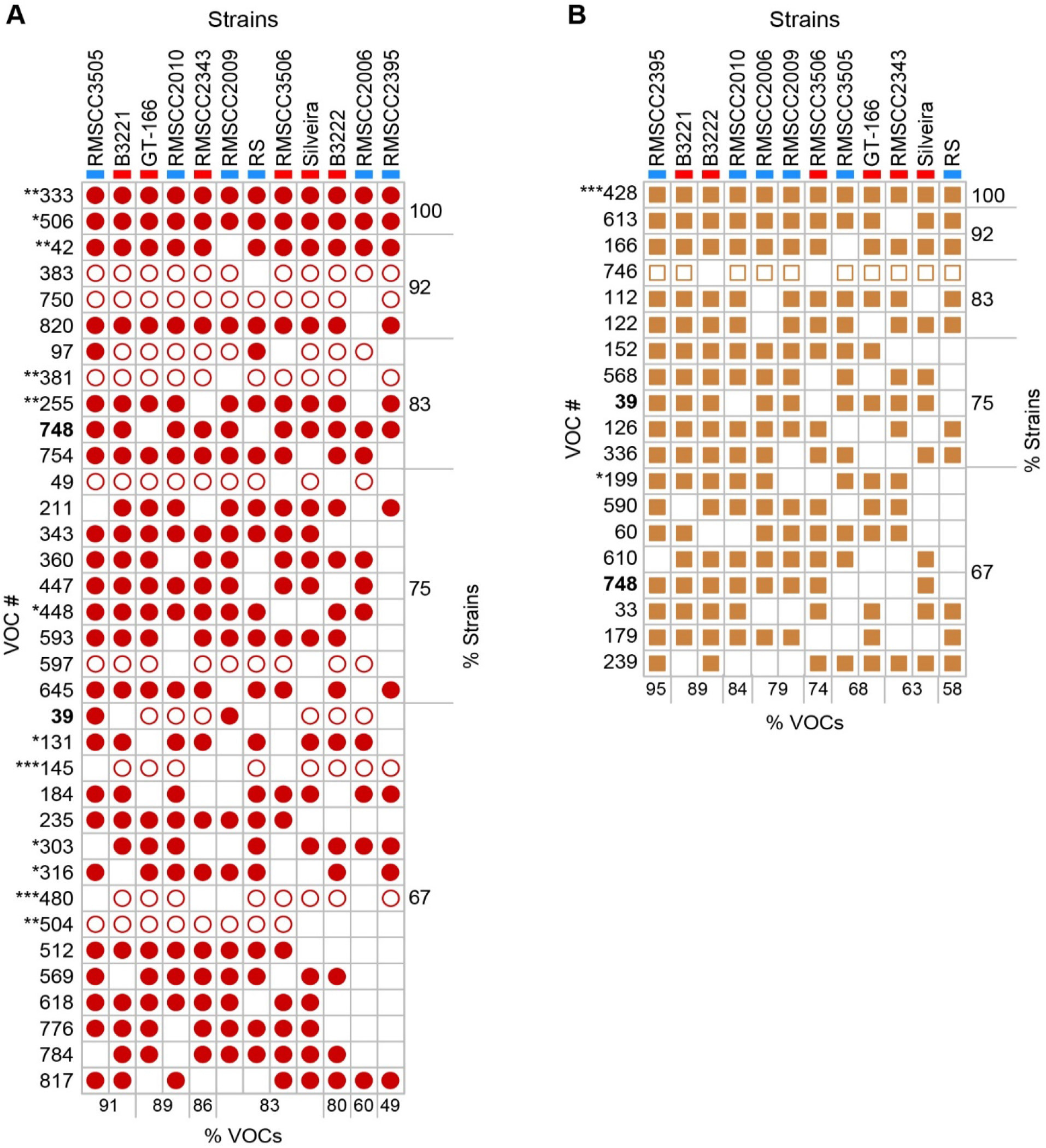
Volatiles detected in at least two-thirds of all *Coccidioides* strains in spherule (A) or mycelial (B) cultures, with VOCs in bold detected in both lifecycles. Open shapes indicate VOCs that are at least 2-fold greater in relative abundance compared to media blanks, and solid shapes indicate VOCs that are at least 10-fold greater in relative abundance. Volatiles marked with an asterisks indicate statistically-significant differences in abundance between lifecycles (* p < 0.05, ** p < 0.001, *** p < 0.0001). Strains are color-coded as *C. immitis* (blue) and *C. posadasii* (red). The percentages of listed VOCs detected in each strain, and the percentages of strains producing each VOC are noted at the bottom and right side of each chart, respectively. Compound IDs: 97 = acetic acid; 239 = 1-hexanol; 255 = 2-hexen-1-ol; 383 = limonene; 428 = 2,4,6,-trimethyl-pyridine; 820 = 2-butyl-1-octanol.

### Multivariate analyses of the *Coccidioides* volatilome

Principal components analysis (PCA), using the 78 cultures and media blanks as observations and 353 *Coccidioides* VOCs as variables, shows that the volatilomes of individual strains and culture conditions are highly reproducible (Figure S1A), and that they cluster by lifecycle, not by species (Figure 4). This observation is reinforced when we include only mycelia or spherules in the PCA, where no separation between species is observed (Figure S1, B and C), or only include *C. immitis* or *C. posadasii*, where we still observe clear separation by lifecycle (Figure S1, D and E). Evaluating these subsets of the data, some interesting variations between strains emerge. The type strain for *C. immitis* is RS; however, in both mycelial and spherule lifecycles, we observe that its volatile metabolome is quite unique from the other *C. immitis* strains (Figure S1). *C. posadasii* also has a metabolic outlier – RMSCC3506 – but only during mycelial growth (Figure S1, B and E). These outliers cannot be fully explained by the size of the volatilomes as being abnormally large or small (Table S2).

**Figure 4.**
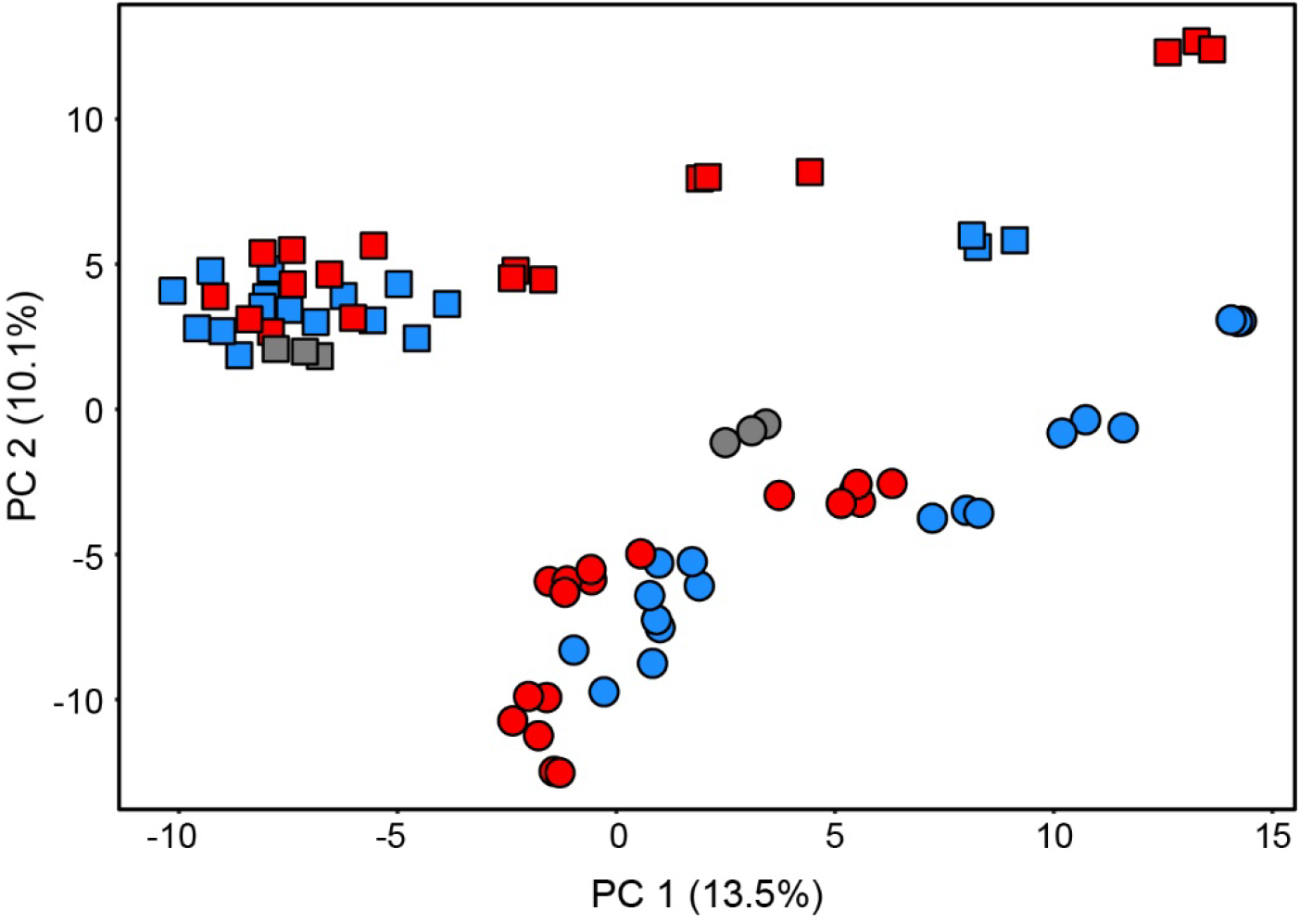
Principal components analysis (PCA) score plot using 353 VOCs as features, produced by *C. immitis* (blue) and *C. posadasii* (red) when cultured in conditions that induced spherule (circle) or mycelial (square) morphologies. Blank media are shown in gray. Fungal cultures and media blanks were analyzed in triplicate, yielding 72 fungal and 6 blank observations. PCA score plots with the observations colored by strain are in Figure S1.

Though lifecycle and strain differences in the *Coccidioides* volatilome are observable in the PCA, less than a quarter of the total variance is captured by the first two PCs. Thus, we also used hierarchical clustering analysis (HCA) to compare the *Coccidioides* volatilomes (Figure 5). The HCA reinforces our finding that lifecycle is the most significant contributor to the *Coccidioides* volatilome, and there are not global similarities in the volatilomes within each species, or within populations of the species. There are two main clusters of VOCs driving the separation of the mycelia and spherule lifecycles in the HCA. Cluster 1 includes 56 VOCs, which are generally present in a higher abundance in mycelia samples compared to spherule samples, five of which are significantly different between lifecycles (p < 0.05 with Benjamini-Hochberg FDR correction; Table S1). In Cluster 1, seven compounds are identified: 3-undecene, 1-ethyl-4-methyl-benzene, 1-methyl-3-(1-methylethyl)-benzene, 2-butyl,1-octanol, decanal, limonene, and β-pinene. As in the PCA, we find that *C. immitis* RS and *C. posadasii* RMSCC3506 are clustering separately when grown as mycelia, partially driven by low abundances of VOCs from Cluster 1. Cluster 2 includes 16 VOCs that are present in spherule samples while generally absent or at a lower abundance in mycelia samples, and all of which are significantly different between lifecycles (p < 0.05 with Benjamini-Hochberg FDR correction). Cyclohexanone and 2,4,6-trimethlpryidine are two VOCs in this cluster that we were able to identify.

**Figure 5.**
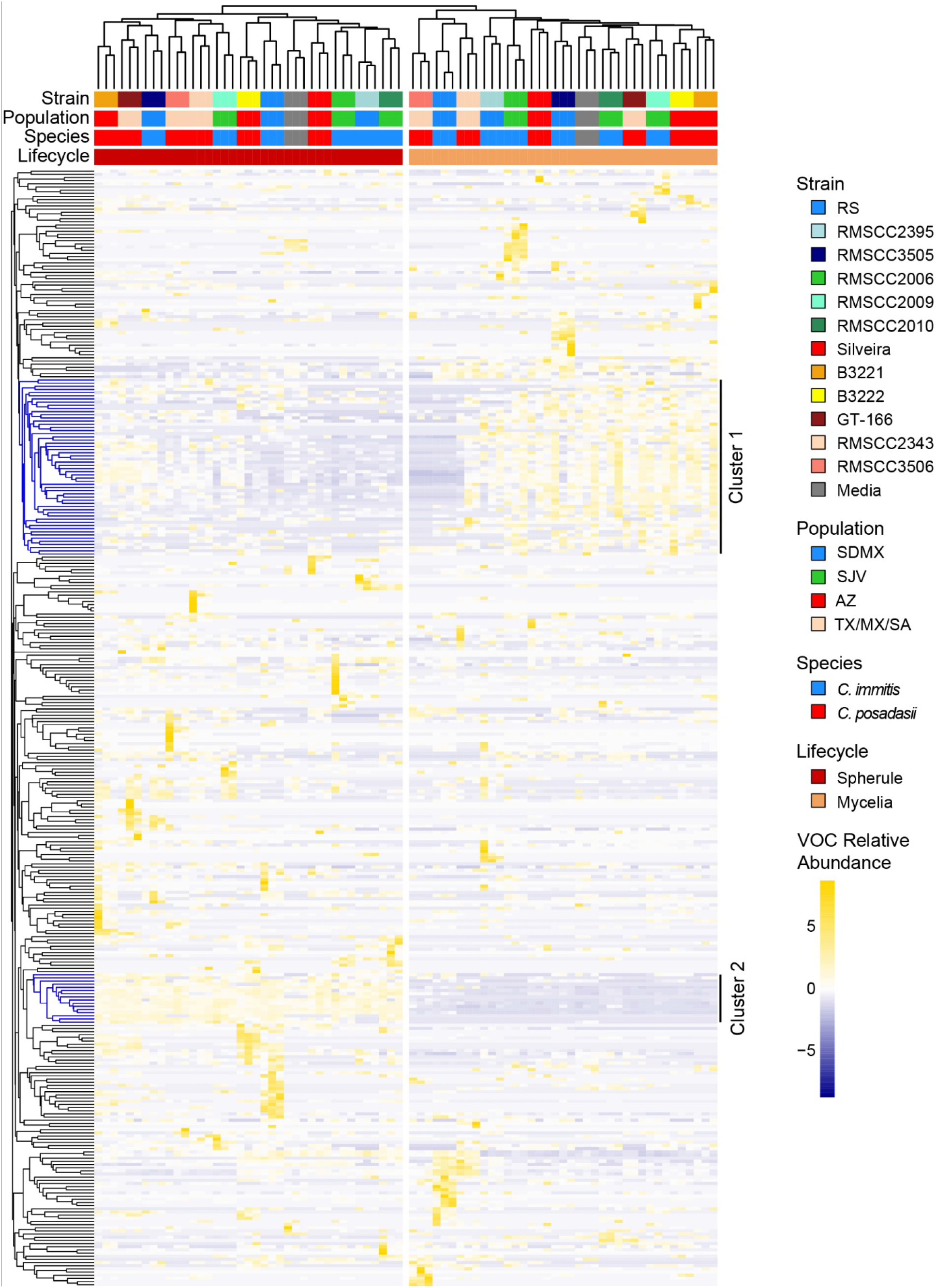
Hierarchical clustering analysis (HCA) of 72 fungal cultures and 6 media blanks (columns) based on the relative abundance of 353 volatiles (rows). Clustering of fungi and blank samples is using Pearson correlations, and clustering of volatiles is using Euclidean distance, both with average linkage. Samples are color-coded by strain, population, species, and lifecycle, as labeled in the legend; media blanks are in gray.

## DISCUSSION

The genus *Coccidioides* is divided into two closely related and putatively allopatric species, with a strong phylogeographic signal and population substructure within each species (19). However, we find that the character of the *Coccidioides* volatilome is determined by lifecycle rather than species. Within each species, the two lifecycles have a relatively small overlap in their shared volatilome, with 27% and 31% shared for *C. posadasii* and *C. immitis,* respectively (Figure 1). While there were no volatiles that were significantly different in relative abundance between *C. posadasii* and *C. immitis*, we observed 43 statistically-significant differences in the VOCs produced during the two lifecycles (Figure 2). This global difference between lifecycles is the primary driver of separation we observed in multivariate analyses using PCA (Figure 4) and HCA (Figure 5). These findings are in line with the large-scale transcriptional changes that occur when *Coccidioides* spp. transitions from mycelial to spherule growth; between 20% and 40% of the *Coccidioides* genes are differentially regulated during the shift between the environmental and parasitic lifecycles (43–46). It has also been shown that the proteomes of the two lifecycles are distinct, with statistically-significant differences between mycelia and spherule lifecycles (47). In contrast to the significant differences in the volatilomes between lifecycles, we observed no uniform differences between the volatilomes of *C. posadasii* and *C. immitis*. There is a high degree of overlap (56 – 57%) between the species’ volatilomes when comparing the same lifecycle (Figure 1). This similarity is also visible in the PCA (Figures 4 and S1) and HCA (Figure 5), as the volatilomes do not cluster by species even when we control for lifecycle. Global similarities between *C. posadasii* and *C. immitis* have also been demonstrated at the protein level (47, 48).

The lack of significant differences between the *C. posadasii* and *C. immitis* volatilomes is due, in part, to large differences observed between strains of the same species; the volatilome highlights some unique aspects of a few of the individual *Coccidioides* strains that we analyzed. For example, RS is the type strain for *C. immitis*; however, it appears to be a metabolic outlier during both lifecycles (Figure S1), but especially during mycelial growth (Figure 5). Additionally, the mycelia lifecycle of *C. posadasii* RMSCC3506 appears to be a metabolic outlier compared to the other strains grown as mycelia (Figures S1 and 5). These mycelia outliers appear to be driven, at least in part, by the lack of volatiles that make up Cluster 1 in the HCA (Figure 5). None of the 56 VOCs found in Cluster 1, which are more abundant in mycelia compared to spherule cultures, are detected in RMSCC3506 cultures, and only two are detected in RS mycelia cultures. These two strains are also lacking VOCs 39 and 748, which were detected in the majority of strains for both lifecycles, but absent from both lifecycles of RS and mycelia cultures of RMSCC3506 (Figure 3). Additionally, none of the 11 VOCs significantly more abundant in mycelia over spherule cultures (Figure 2 and Table S1) are detected in RMSCC3506, and only two are detected in RS. Beyond contributing to high levels of volatilome variance in *Coccidioides*, the significance of the strain level differences relative to fungal biology or infection are not known. Variation in virulence in murine models of coccidioidomycosis among strains has been observed, but has not been tested in the context of species or with a mechanistic hypothesis to test (49). In future studies of mouse model and human infections we plan to investigate correlations between the *Coccidioides* volatilome and virulence and infection outcomes.

It should be noted that although the spherule and mycelia media had the same composition, we detected different VOCs in the headspace of the blanks (Table S1, Figures 4 and 5). We interpret these differences as the influence of the cultures on the blanks, rather than the influence of the blank media on the cultures. Because the media blanks were incubated and processed in parallel with their respective samples, we posit that the blank media absorbed volatiles from the culture environment (e.g., from fungal spherule or mycelial cultures in shared incubators), which introduced differences in their volatilomes. An analysis of fresh blank media (not incubated with the cultures) would be required to test this hypothesis. Our data indicate that the spherule media absorbed a greater abundance of compounds compared to the mycelia media (Table S1), which may be due to the spherule lifecycle producing greater numbers and/or abundances of individual compounds. Therefore, it is possible there are additional significant spherule volatiles that were not identified within this study, since a VOC was defined as “detected” based on its relative abundance versus media blanks. Future *in vitro* work needs to be mindful of volatile absorption by the cultures and blanks within a shared space, such as an incubator.

The results from this study highlight two crucial factors that need to be considered in the development and translation of volatilomics (and other functional genomics data) into biomarkers. First, there can be a significant amount of intra-species variation, such that the volatilome of a single strain is not representative of the species as a whole (17, 27, 50, 51). Therefore, the analysis of multiple strains should be performed to identify a core volatilome, and the more genetically and phenotypically diverse the species, the more strains will need to be analyzed to determine the consensus set of volatiles produced by the majority of them. Second, our data show that the growth phase of the organism has a strong influence on the volatilome, and therefore using *in vitro* culture conditions that can most closely replicate the *in vivo* environment and microbial growth phases may increase the likelihood of putative biomarker translation from bench to bedside. In addition to controlling the growth phase of dimorphic organisms, macronutrient, micronutrient, and oxygen availability should all be considered in the development of *in vitro* model systems that replicate *in vivo* physiology of the pathogen (14, 17, 27, 30, 33, 36, 52, 53). Our next step in developing a Valley fever breath test is to translate *in vitro* biomarkers for use with an *in vivo* mouse model. We observed 35 VOCs present in at least 67% of all spherule strains (Figure 3); as this is the parasitic lifecycle found within the host, we hypothesize that the most highly conserved spherule VOCs have the greatest probability of translating to Valley fever breath biomarkers.

## MATERIALS AND METHODS

### Fungal Isolates and Growth Conditions

#### Isolates

The *Coccidioides* isolates used in the study were obtained from human patients and are listed in Table 1. All *Coccidioides* isolates were grown under BSL-3 containment, using conditions that induce mycelial or spherule growth.

For mycelial growth, a 50 ml vented falcon tube containing 10 ml of RPMI media (filter sterilized RPMI 1640, 10% fetal bovine serum) was inoculated with a 1 cm × 1 cm 2xGYE agar plug for each strain. These plates were inoculated using 100 μl of glycerol stock, spread across the plate, and cultured for 30°C for two weeks. Control RPMI media was inoculated with a plug from sterile 2xGYE agar media. Each sample, including media control, was prepared in triplicate. Cultures were grown on a shaking incubator at 150 rpm, 30°C for 96 h. For spherule cultures, a 50 ml vented falcon tube containing 10 ml of RPMI media was inoculated to a final concentration of 1.0 × 10^5^ arthroconidia/ml in 1x phosphate-buffered saline (PBS). Arthroconidia were grown and harvested as previously described (54). Strains RMSCC2343 and RMSCC3505 did not produce enough conidia to achieve 1.0 × 10^5^ arthroconidia/ml, and were inoculated at 7.0 × 10^4^ and 4.0 × 10^4^ arthroconidia/ml, respectively. Control media was inoculated with 1 ml of sterile 1xPBS. Cultures were grown on a shaking incubator at 150 rpm, at 39°C in 10% CO_2_ for 96 h. Mycelial and spherule cultures were spun at 12,000 × g at 4°C for 10 min to pellet the cells. The supernatant was removed and place in a Nanosep MF Centrifugal Devices with Bio-Inert® Membrane 0.2 μm spin filter and centrifuged at 3,200 × g for 4 min. The filtrate was stored at −80°C until volatile metabolomics analysis.

To ensure sterility of the filtrates for metabolomics analyses outside of a BSL-3 containment facility, 10% of all sample filtrates were plated on 2xGYE and incubated at 30°C for 96 h to ensure complete removal of viable pathogen particles. No growth was observed in any replicate.

### Volatile metabolomics analysis by SPME-GC×GC–TOFMS

The *Coccidioides* spp. culture filtrates and media blanks were allowed to thaw at 4°C overnight, and then 2 ml were transferred and sealed into sterilized 10 ml GC headspace vials with PTFE/silicone septum screw caps. All samples were stored for up to 12 d at 4°C until analyzed. Samples were randomized for analysis. Volatile metabolites sampling was performed by solid-phase microextraction (SPME) using a Gerstel^®^ Multipurpose Sampler directed by Maestro^®^ software. Volatile metabolite analysis was performed by two-dimensional gas chromatography−time-of-flight mass spectrometry (GC×GC–TOFMS) using a LECO Pegasus 4D and Agilent 7890 GC (St. Joseph, MI). Chromatographic, mass spectrometric, and peak detection parameters are provided in Table S3 (GC parameters). An external alkane standards mixture (C_8_ – C_20_; Sigma-Aldrich, St. Louis, MO) was sampled multiple times for calculating retention indices (RI). The injection, chromatographic, and mass spectrometric methods for analyzing the alkane standards were the same as for the samples.

### Processing and analysis of chromatographic data

Data collection, processing, and alignment were performed using ChromaTOF software version 4.71 with the Statistical Compare package (Leco Corp.), using the parameters listed in Table S3 (GC parameters).

Peaks were assigned a putative identification based on mass spectral similarity and RI data, and the confidence of those identifications are indicated by assigning levels 1 – 4 (1 highest) (55). Peaks with a level 1 ID were identified based on mass spectral and retention index matches with external standards. Peaks with a level 2 ID were identified based on ≥ 800 mass spectral match by a forward search of the NIST 2011 library and RI that are consistent with the midpolar Rxi-624Sil stationary phase, as previously described (50), but using an RI range of 0 – 43% (empirically determined by comparing the Rxi-624Sil RIs for Grob mix standards to published polar and non-polar values). Level 1, 2, and 3 compounds were assigned to chemical functional groups based upon characteristic mass spectral fragmentation patterns and second dimension retention times, as previously described (51). Level 4 compounds have mass spectral matches < 600 or retention indices that do not match previously published values, and are reported as unknowns.

### Data post processing and statistical analyses

The data post processing steps are depicted in Figure S2. Before statistical analyses, compounds eluting prior to 358 s (acetone retention time) and siloxanes (*i.e.,* chromatographic artifacts) were removed from the peak table. Peaks that were present in only one of the three biological replicates were imputed to zero for that sample, while missing values for peaks that were present in two out of three biological replicates were imputed to half of the minimum value across all biological replicates. The relative abundances of compounds across chromatograms were normalized using probabilistic quotient normalization (PQN) (56) in R version 3.4.3. The data were log10 transformed, and intraclass correlation coefficients (ICCs) were calculated using R *ICC* package version 2.3.0, and peaks with an ICC < 0.75 were not further processed. Analytes were retained for further analysis if they were at least two-fold more abundant in any *Coccidioides* culture compared to media controls, and were considered to have been detected in that culture. Principal component analysis was performed using R *factoextra* package version 1.0.5 with the biological replicates as observations and the absolute peak intensities (mean-centered and scaled to unit variance) as variables. The relatedness of samples based on their volatile metabolomes were assessed using hierarchical clustering analysis on the Pearson’s correlation between isolates and Euclidean distance between volatiles using R *pheatmap* package version 1.0.12. Geometric means of the biological replicates were calculated and used to determine the statistical difference in abundance between lifecycles or species using a Mann-Whitney *U*-test and Benjamini and Hochberg false discovery rate correction with α = 0.05, using R *stats* package version 3.5.3. The variation in abundance of volatiles significantly different between lifecycle was calculated by dividing mean mycelia peak abundance by mean spherule peak abundance.

### Data availability

Metabolomic data (chemical feature peak areas and retention time information) included in this study are available at the NIH Common Fund’s National Metabolomics Data Repository (NMDR) website, the Metabolomics Workbench, at www.metabolomicsworkbench.org, where it has been assigned project ID PR000### and study ID ST00### (https://doi.org/####).

## CONCLUSION

The *Coccidioides* volatilome is strongly influenced by the environmental versus parasitic lifecycles, while the two species, *C. posadasii* and *C. immitis*, are indistinguishable by their volatile profiles, in part due to high strain to strain variability within the species. We identified 35 VOCs produced by the majority of the *Coccidioides* strains when grown as spherules, the parasitic form of the fungus found in Valley fever lung infections. Our results show that as future efforts are made to develop biomarkers for *Coccidioides* and other mycoses caused by dimorphic fungi, it is vitally important to control for fungal lifecycle and sample multiple strains of the pathogen to identify volatiles that are more highly conserved within the infectious genus or species.

## ACKNOWLEDGEMENTS

Funding for this work was provided by the Arizona Biomedical Research Commission (ABRC) award ADHS18-198861 (HDB). We thank John Galgiani and George Thompson who provided *Coccidioides* clinical isolates to BMB.

We declare that the research was conducted in the absence of any commercial or financial relationships that could be construed as a potential conflict of interest.

