## Supplemental Figures and Tables for "Lifecycle dominates the volatilome character of the dimorphic fungus *Coccidioides* spp"

Running Title: Lifecycle dominates the volatilome of *Coccidioides*

#### Results

**Table S1.** See Supplementary Excel File. Table of 353 *Coccidioides* VOCs detected in the headspace of *in vitro* spherule and mycelial cultures.

**Table S2.** The numbers of volatiles detected in at least two-fold greater concentration in *Coccidioides* cultures versus blank media. The type strains for each species are indicated with an asterisk.

| **Strain** | **Mycelia (n = 224)** | **Spherule (n = 272)** |
|---|---|---|
| ***C. posadasii* (n = 291)** | | |
| Silveira* | 51 | 61 |
| B3221 | 56 | 122 |
| B3222 | 55 | 80 |
| RMSCC2343 | 69 | 80 |
| RMSCC3506 | 51 | 77 |
| GT-166 | 36 | 107 |
| ***C. immitis* (n = 309)** | | |
| RS* | 71 | 105 |
| RMSCC2395 | 81 | 42 |
| RMSCC3505 | 63 | 96 |
| RMSCC2006 | 59 | 53 |
| RMSCC2009 | 54 | 68 |
| RMSCC2010 | 61 | 75 |

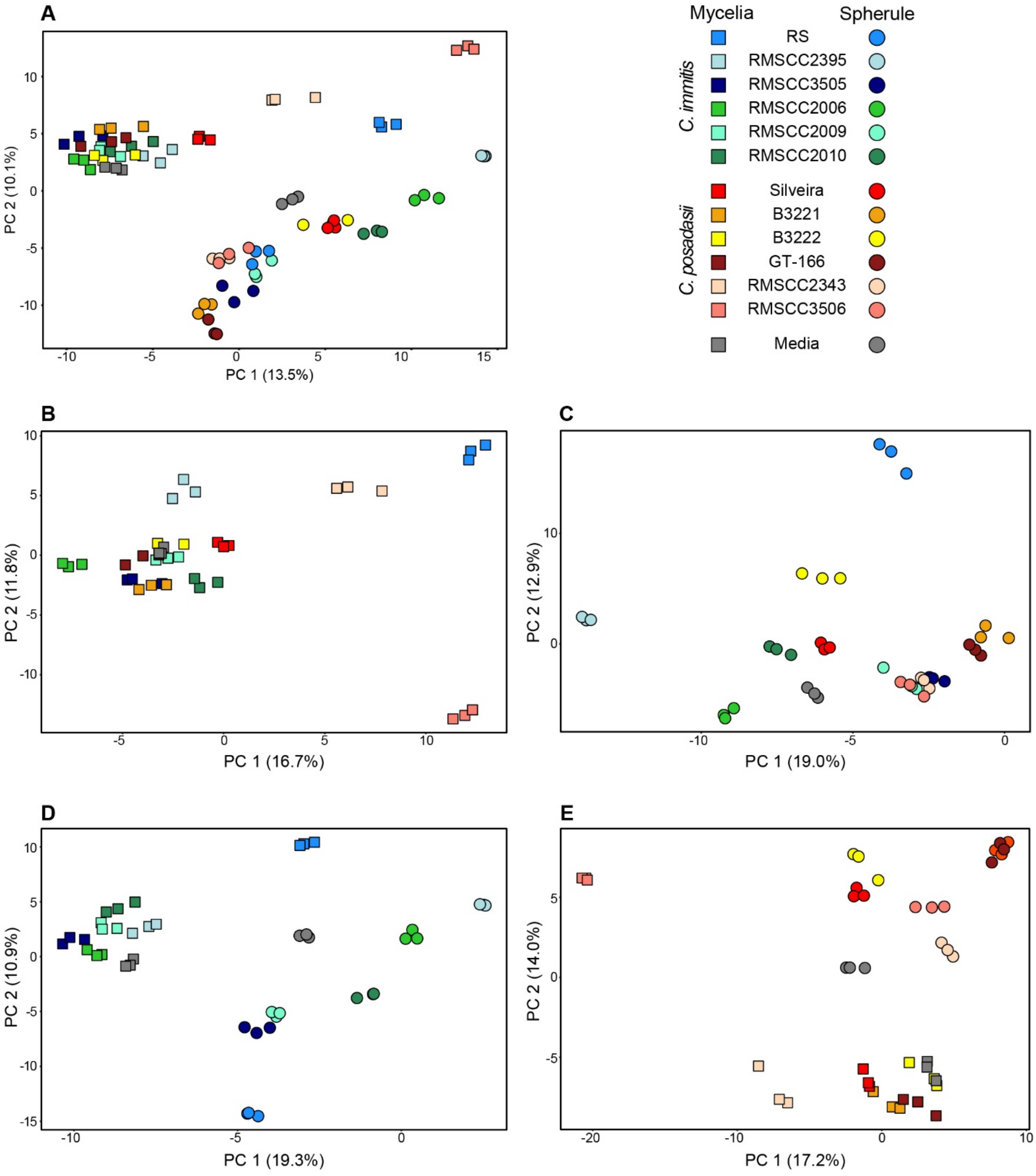


**Figure S1.** Principal components analysis (PCA) score plot using VOCs as features, produced by 12 strains of *C. immitis* and *C. posadasii* when cultured in conditions that induced mycelial (square) or spherule (circle) morphologies. Fungal cultures and media blanks were analyzed in triplicate. Fungal strains and the media blanks are color coded, as labeled in the legend. (A) All samples as observations and 353 VOCs as features. (B) Mycelial cultures and blanks as observations and 224 VOCs as features. (C) Spherule cultures and blanks as observations and

272 VOCs as features. (D) *C. immitis* cultures and blanks as observations and 309 VOCs as features. (E) *C. posadasii* cultures and blanks as observations and 291 VOCs as features.

**Methods**

**Table S3.** Parameters for HS-SPME and GC×GC-TOFMS analysis, and data processing and alignment

| | |
|---|---|
| **Autosampler Method** | |
| Instrument description | Gerstel® MPS Pro® |
| Software description | Gerstel® Maestro® (version 1.5.3.2) |
| **Sampling Parameters** | |
| Cooled tray temperature | 4 °C |
| Solid-phase microextraction (SPME) | Manufacturer: Supelco®<br>Fiber type: PDMS/CAR/DVB (1 cm; 50/30 µm) |
| Incubation time | 5 min |
| Agitator parameters, incubation | Temperature: 50 °C<br>On time: 10 s<br>Off time: 1 s<br>Speed: 600 rpm |
| Agitation, sampling | On |
| Vial penetration | 21 mm |
| Extraction time | 10 min |
| Injection penetration | 67 mm |
| Desorption time | 180 s |
| **Inlet (CIS) Parameters** | |
| Initial temperature | 250 °C |
| Equilibrium time | 0.05 min |
| Initial time | 0.10 min |
| Ramp rate | 12 $°C \cdot s^{-1}$ |
| End temperature | 250 °C |
| Hold time | 10.5 min |
| **GC×GC Method** | |
| Instrument description | Agilent® 7890B |
| Column configuration | Column 1: Rxi®-624Sil MS, 60 m × 0.25 mm × 1.4 $\mu$m<br>Column 2: Stabilwax®, 1 m × 0.25 mm × 0.5 $\mu$m |
| Carrier gas | Helium, 2 $mL \cdot min^{-1}$ (constant) |
| Front inlet type | Gerstel® |
| Front inlet mode | Splitless |
| Front inlet septum purge flow | 1 $mL \cdot min^{-1}$ |
| Front inlet septum purge time | 300 s |

| | |
|---|---|
| Front inlet purge flow | 50 mL·min$^{-1}$ |
| Front inlet total purge flow | 52 mL·min$^{-1}$ |
| Oven equilibration time | 5 s |
| Primary oven temperature ramp | Initial temperature: 35 °C<br>Initial time: 0.5 min<br>Ramp rate: 5 C·min$^{-1}$<br>Final temperature: 230 °C<br>Hold time: 5 min |
| Secondary oven temperature offset | +5 °C (relative to primary oven) |
| Modulator temperature offset | +15 °C (relative to secondary oven) |
| Modulation timing | Modulation period: 2.00 s<br>Hot pulse time: 0.50 s<br>Cold pulse time: 0.50 s |
| Transfer line temperature | 250 °C |

| Mass Spectrometry Method | |
|---|---|
| Instrument description | LECO® Pegasus® 4D |
| Use GC method total time for MS method total time | Yes |
| Acquisition delay | 180 s |
| Filament active time | 180 s to end of run |
| Start mass/End mass | 35/400 |
| Acquisition rate | 100 spectra·s$^{-1}$ |
| Optimized voltage offset | +50 V |
| Electron energy | -70 eV |
| Ion source temperature | 250 °C |

| Data Processing Method | |
|---|---|
| Software description | LECO® ChromaTOF® and<br>Statistical Compare (version<br>4.71.0.0) |
| Baseline tracking/Offset | Entire run/0.5 (through middle of noise) |
| Data points averaged for smoothing | Auto |
| First dimension peak width | 12 slices |
| Mass spectral match required to combine | 600 |
| Second dimension peak width | 0.15 |
| Min. subpeak signal-to-noise (S/N) for | 6 |
| Integration approach | Traditional |
| Peak finding | S/N: 50<br>Number of apexing masses: 2 |
| Mass spec libraries for searching | NIST 2011 |
| Mass to use for area/height calculation | Unique mass |
| Alignment analyte match criteria | Spectral match mass threshold: 10<br>Minimum spectral similarity match: 600<br>Max. number of modulation periods apart: 3<br>Max. retention time difference (s): 0.2<br>S/N for second peak find: 5 |
| Criteria for inclusion of analytes | Min. number of samples that contain analyte: 1 |

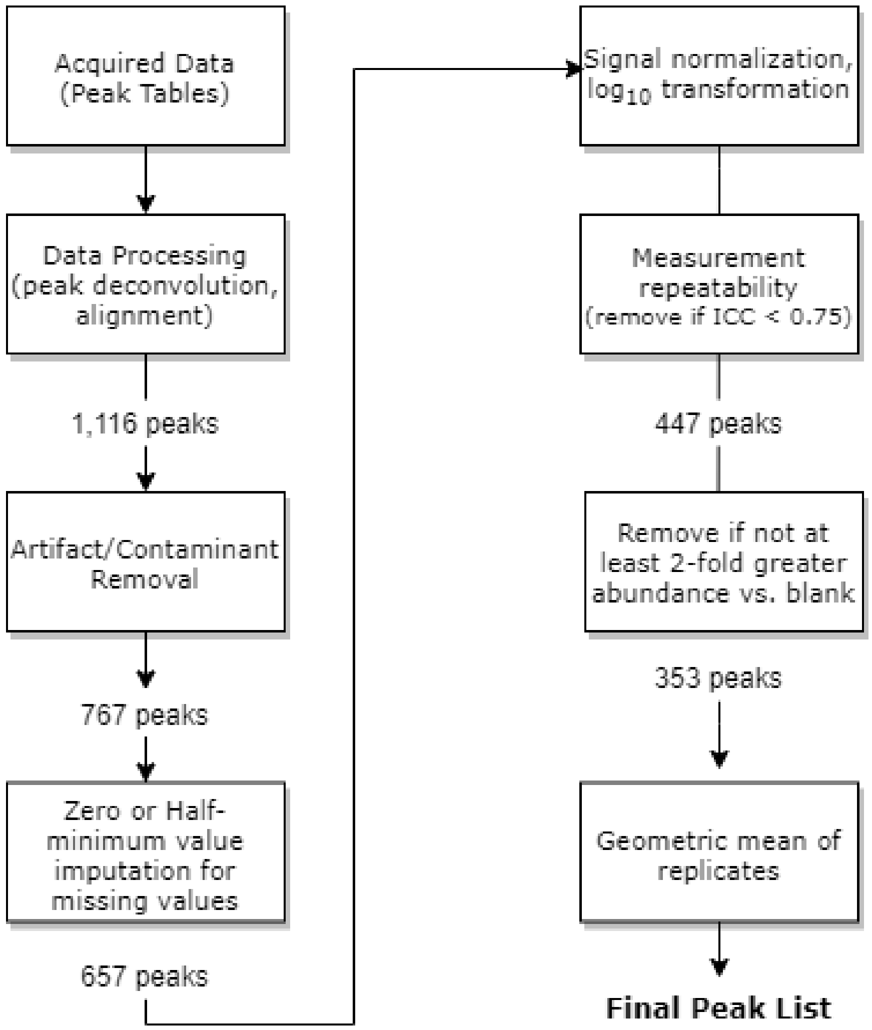


**Figure S2.** Data post processing workflow
